# Cooperation & Liaison between Universities & Editors (CLUE): Recommendations on Best Practice: Preprint for Consultation

**DOI:** 10.1101/139170

**Authors:** CLUE Working Group, Elizabeth Wager, Sabine Kleinert, Michele Garfinkel, Volker Bahr, Ksenija Bazdaric, Michael Farthing, Chris Graf, Zoë Hammatt, Lyn Horn, Susan King, Debra Parrish, Bernd Pulverer, Paul Taylor, Gerrit van Meer

## Abstract

Journals and research institutions have common interests regarding the trustworthiness of research publications but their specific roles and responsibilities differ. These draft recommendations aim to address issues surrounding cooperation and liaison between journals and institutions about possible and actual problems with reported research. The proposals will be discussed at various meetings including the World Conference on Research Integrity in May 2017. We will also consider comments and suggestions posted on this preprint.

The main recommendations are that:

- National registers of individuals or departments responsible for research integrity at institutions should be created.
- Institutions should develop mechanisms for assessing the validity of research reports that are independent from processes to determine whether individual researchers have committed misconduct.
- Essential research data and peer review records should be retained for at least 10 years.
- While journals should normally raise concerns with authors in the first instance, they also need criteria to determine when to contact the institution before, or at the same time as, alerting the authors in cases of suspected data fabrication or falsification to prevent the destruction of evidence.
- Anonymous or pseudonymous allegations made to journals or institutions should be judged on their merit and not dismissed automatically.
- Institutions should release relevant sections of reports of research trustworthiness or misconduct investigations to all journals that have published research that was the subject of the investigation.

## Background

Journals and research institutions (e.g. universities) share common interests when concerns arise over the trustworthiness of research reports that are submitted for publication or published. This shared interest means that cooperation, in the form of sharing information, is often necessary. Concerns about the reliability of reported research may arise during editorial assessment or peer review or from pre-publication screening (e.g. for plagiarism or image manipulation) therefore journals may be the first to suspect problems. However, journals usually do not have all the evidence, or a specific mandate, to conduct a formal investigation. Therefore it is important for them to alert the relevant institution(s) and funder(s). Liaison between institutions and journals is also important after an institutional investigation, especially if the investigation indicates that published work may not be reliable (for whatever reason), so that the research record can be corrected. However, cooperation between journals and research institutions is not always straightforward and both report difficulties and frustrations.

In 2012, the Committee on Publication Ethics (COPE) published guidelines on cooperation between research institutions and journals on research integrity cases [1]. These guidelines were discussed at the World Conference on Research Integrity in Montreal in 2013 and a series of questions was formulated on which further guidance was desired [2]. This document is largely based on those questions.

## Development of this document

In July 2016, a meeting was held in Heidelberg, hosted by the European Molecular Biology Organization (EMBO) with financial support from COPE. The aim of the meeting was to address the questions raised in Montreal, to understand the reasons why communication and cooperation between journals and institutions is sometimes challenging, and to identify practical solutions to problems. The meeting brought together editors and publishers of scholarly journals, people working at universities and national research integrity organizations (including research integrity officers, a university vice-chancellor and a dean), a lawyer with experience of representing researchers, journals and universities in research misconduct cases, and policy experts. The participants came from Australia, Croatia, Germany, the Netherlands, South Africa, UK, and USA.

## Scope

These recommendations cover interactions between representatives of scholarly, peer-reviewed journals and research institutions about cases in which there are concerns about the trustworthiness, honesty, integrity or attribution of reported research that has been submitted for publication to the journal whether or not it has been (or will be) published.

## Terminology

The term “journal” refers to editors and publishing staff who handle cases or develop policy on research and publication integrity. The acronym CLUE (standing for Cooperation and Liaison between Universities and Editors) uses the term “universities” to include all types of research institution (mainly focusing on academic institutions) and “editors” to refer to all journal representatives.

This document does not attempt to define or limit types of research or publication misconduct. During discussion, it was agreed that focusing on narrow definitions of misconduct contributes to the difficulties that sometimes hamper communication between journals and research institutions. As noted in the COPE guidelines, journals have responsibility for the trustworthiness (or soundness) of what they publish and this does not always align with institutions’ definitions of research misconduct [1]. In other words, it is possible for research reports to be misleading or untrustworthy and therefore to require correction or retraction even when the authors/researchers are not considered to have committed research misconduct by their institution.

In this document, therefore, the term “misconduct” is used to describe any actions of researchers that result in research that cannot be trusted, is not reliable, is not presented honestly, and, for whatever reason, should not become part of, or remain on, the research record. It is not based on any particular definition of research misconduct.

The terms “inquiry” and “investigation” refer to formal processes conducted by research institutions to determine whether a researcher/employee has committed misconduct. One of the issues discussed at the CLUE meeting was the extent to which journals should assess evidence of misconduct. While it was agreed that it is not usually the role of journals to conduct formal research misconduct investigations, we recognise that, in some cases, it may be appropriate for journals to consider evidence relating to the integrity of a publication or submission. Institutional investigations tend to focus on the guilt or otherwise of the researcher(s) concerned and seek to determine whether their behaviour amounts to research misconduct however that is defined. However journals are more concerned with whether the research can be trusted and is properly reported and reliable. These are different questions that are answered in different ways and carry different obligations. Journals may conduct their own assessments of the integrity of the research reported in a manuscript or article, but such assessments are often limited by the access that the journal has to all of the necessary information. Institutional consideration may centre more on the behaviour or motivations of the researcher(s) but may not fully address the questions of trustworthiness or reliability that the journal needs to be answered.

## Recommendations on best practice

### Issue: Journals often have difficulty identifying somebody responsible for research integrity at an institution

Journals often report difficulties in identifying the appropriate person to contact at an institution to raise concerns about research integrity. The situation varies by country, but in many areas, universities either do not have a research integrity office or officer (RIO), or the person or department with responsibility for research integrity (and their contact details) are not clearly identified on the institution’s website. Identifying the right contact person is also difficult because different titles are used for this function.

#### Recommendations

Institutions should have a research integrity officer (or office) and publish their contact details. National research integrity bodies (or other appropriate organizations, e.g. major funders) should keep a register of people responsible for research integrity at their country’s institutions, to enable journal editors (and others) to contact them.

Where such lists are not available, journals should request corresponding authors to provide the name and email address (or telephone number) of their institution’s RIO (or of an individual with responsibility for handling research integrity cases).

Note: If the corresponding author’s institution does not have a RIO, the authors may identify a suitable person at any of their institutions. If no such person can be identified at any of the institutions involved with the research, the authors should be asked to nominate a senior faculty member (e.g. dean or pro-vice chancellor with responsibility for research, or the chair of the research ethics committee or institutional review board) who was not directly involved with the research (and is not an author) who could be contacted if the journal has any concerns about research integrity.

Requiring researchers to provide contact details of a person with responsibility for research integrity at their institution should not only enable journals to contact this person if concerns arise, but may also encourage institutions to make such an appointment, raise awareness of RIOs among researchers, and publish their contact details prominently on institutional websites. Details of the contact person for research integrity enquiries should not be published by the journal, but should be retained, should the need arise to contact them.

### Issue: Journals do not know the best way to contact an institution and whether an informal “off the record” discussion is possible

Since journals are typically not in a position (either legally or practically) to conduct formal investigations into misconduct it is not always possible for journals to obtain clear evidence or to judge whether an allegation is well-founded on the basis of submitted or published work. While journals may request source data from authors, they do not have legal powers to obtain this, nor do they have access to laboratory notebooks or equipment logs, or the possibility to interview staff. Therefore, since journals do not normally have access to all the relevant information, their peer reviewers and editors may only be able to indicate they suspect that something is wrong, without being able to define the problem precisely.

Therefore, journals sometimes want to contact institutions informally, to discuss their suspicions or concerns, or raise non-specific allegations, without necessarily invoking a full investigation. Journals may also wish to know whether a researcher is currently being, or has recently been, investigated for suspected misconduct.

Journals need to understand that in some jurisdictions (for example, the United States) such an “off the record” discussion is not always possible, as institutional research integrity officers and all those involved with investigations have to maintain the maximum confidentiality possible until an inquiry has concluded and such conversations must be documented as part of the institutional record. Institutions risk being sued if they breach this confidentiality, e.g. by revealing that a researcher is under investigation.

However, in other regions, the situation is different and it may be possible to discuss concerns informally and for universities to disclose whether an individual is currently under investigation.

### Issue: Should journals always contact authors about research integrity concerns?

In most cases, when journals have concerns about the reliability or integrity of submitted or published work, they should first raise them with the authors (usually starting with the corresponding author). This allows researchers to provide clarification, explanation or further information. Contacting authors is considered to reflect “due process” or procedural fairness, and avoids wasting institutional and editorial time and resources over issues that arise from honest error and that can be handled in a straightforward way by the journal. When approaching authors, journals are advised to describe concerns using neutral rather than accusatory language, for example highlighting the amount of text similarity rather than accusing an author of plagiarism. The presumption, at this stage, is that the authors are “innocent until proven guilty”.

However, journals should be aware that in cases of suspected data fabrication or falsification, raising concerns with the authors first could enable researchers to destroy or alter evidence that might be important for an institutional investigation [3,4]. Therefore, when journals have well-founded suspicions or evidence of falsification or fabrication they should consider informing the institution at the same time as, or before, they contact the author(s).

Such cases are likely to be rare, since the circumstances in which journals have access to raw data are currently limited (but may include western blots and other images). This situation may change as publication of research data becomes more widespread [5].

If a journal discovers evidence of falsification (e.g. inappropriate manipulation of images) or major plagiarism (e.g. reports from text-matching software verified by an editor) the journal should retain the evidence and should offer to share it with the institution. However, care should be taken to avoid revealing the identity of peer reviewers, or other people raising concerns, to an institution against their wishes or without their permission. Ensuring the anonymity of internal whistleblowers (i.e. members of a research group or department who raise concerns about colleagues or collaborators) may be difficult since, even if their name is not revealed, the source may be obvious to the authors if only a few people would know about certain details of the research.

#### Recommendation

Journals should develop criteria to determine when the authors’ institution(s) should be contacted immediately without (or at the same time as) alerting the author(s). (See, for example the EMBO Press classification for image aberrations [6].) This would normally occur only in exceptional cases when journals have strong suspicions or clear evidence of substantive or significant falsification or fabrication of data.

### Issue: What should journals do when reviewers say findings look “too good to be true” in the absence of specific evidence?

If a peer reviewer raises a concern about the trustworthiness of findings, especially if s/he suggests that the results are “too good to be true”, the journal should ask them for more details (e.g. to explain why they gave this opinion) and should usually alert the institution to these concerns if they consider they are well founded. Journals therefore need to determine whether to contact an institution and, if so, what information they should share.

Peer reviewer reports and comments to the editor should generally only be shared with authors’ institutions with the reviewers’ express permission. Similarly, the identity of the peer reviewer should not normally be revealed to the authors’ institutions in cases of suspected problems with a submitted or published work

It is helpful for journals to share suspicions about the reported research with institutions (as well as more specific concerns or clear evidence) because institutions are able, and have a duty, to assess concerns about data fabrication or falsification by researchers. Another reason why journals should raise non-specific concerns about reported research is that the institution should have a more complete picture of the researcher’s behaviour than the journal (which usually has information only from one article), and such evidence may be important to trigger or inform an investigation. Sophisticated data fabrication or falsification may only become obvious when several publications are assessed, or when raw data or other forensic evidence are available [7]. Therefore, in such cases, while individual journals may have some suspicions, the full picture is available only to the institution. Furthermore, alerting the institution may prevent the research from being submitted to other journals (which would be unaware of the first journal’s concerns) before it has been properly assessed.

#### Recommendations

Journals should develop criteria for determining whether, and what type of, information should be passed on to institutions.

Journals should share evidence relating to possible misconduct with institutions but should not reveal the identity of peer reviewers or other people raising concerns (unless this is already published or the individuals have given permission for this disclosure).

In addition to sharing any direct evidence of plagiarism, fabrication or falsification with institutions, journals should share reviewer or editor suspicions that work is “too good to be true” or a strong suspicion of something being “not right”.

### Issue: Investigating the reliability of reported research usually requires access to original (raw) data but these may not be available for research done several years ago

It is often impossible to investigate the validity and integrity of a piece of research and its reporting without access to the raw data. This can be problematic if data are not retained, since suspicions sometimes emerge several years after publication. Also, if data are kept only by individual researchers, files may be lost unintentionally or deliberately destroyed or altered.

Permanent, public deposition of data is the ideal, since it allows immediate scrutiny by anybody interested, which may reveal errors or misconduct. However public posting of individuals’ personal or clinical data may not be possible due to the need for confidentiality (e.g. of medical records).

We encourage institutions and funders to review current data retention standards which may prevent effective investigation of historical data (e.g. we suggest that the 6-year period required for the retention of personal health data in the US under the Health Insurance Portability and Accountability Act (HIPAA) [8,9] is too short). We also encourage debate on the risks and benefits of conventions in certain disciplines of destroying sensitive data, such as interview transcripts, to protect the confidentiality of research participants and to develop alternative systems (e.g. locked, secure deposition) to permit later investigation, if required.

Similarly, investigation of peer review manipulation requires access to journals’ editorial records [10, 11]. Publishers should therefore retain records for a similar period.

#### Recommendations

Research institutions and major funders should have systems to ensure that essential research data are retained for at least 10 years, and ideally permanently. Responsibility for data storage (e.g. for multicentre studies) should be defined in funding agreements.

Journals and publishers should retain peer review records for similar periods to enable the investigation of peer review manipulation or other inappropriate behaviour by authors or reviewers.

### Issue: Institutional focus on strict definitions of research misconduct may hamper communication about broader issues of research integrity and reliability

Journals have a responsibility to correct or retract any publications that give misleading accounts of research methods, findings, analyses or authorship, regardless of whether this is determined to have been due (or related) to misconduc or to error. However, many institutions and research integrity bodies focus solely on determining narrowly defined misconduct and establishing the burden of proof for each particular case. Furthermore, definitions of misconduct vary between jurisdictions. For example, the US Office of Research Integrity considers only cases of fabrication, falsification or plagiarism (FFP) in research funded by the US Public Health Service [12] while the draft Australian code for the responsible conduct of research takes a more inclusive approach [13].

Because of the possible serious consequences of a misconduct finding for individuals and institutions and the importance of conducting rigorous and fair proceedings (and the costs associated with these), thresholds for launching a full inquiry or investigation may be high. This may give journals the impression that institutions are reluctant to cooperate or respond to their enquiries.

It would therefore be helpful if institutions had mechanisms for assessing the validity of reported research in response to concerns raised by journals or others. The focus of such assessment should be solely on determining the trustworthiness of the research itself, and its reporting, rather than on the behaviour or intentions of the researchers. Such assessments should permit institutions to respond more rapidly to journal enquiries and without concerns about breaching confidentiality related to institutional policies or employment processes. However, such assessments would not prevent further investigation through the institution’s established processes for handling misconduct allegations.

#### Recommendation

Institutions should develop mechanisms for assessing the validity of research reports that are submitted to, or published by, academic journals; these should be independent from processes to determine whether misconduct has occurred.

### Issue: Institutions may feel legally bound to keep disciplinary hearings confidential and may therefore feel unable to communicate or share details of on-going investigations with journals

Journals have a responsibility to alert readers to published material that may be untrustworthy. Even when misleading research does not cause direct public harm, it may lead to the waste of other researchers’ time and resources. The need for journals to alert readers promptly to potentially unreliable articles is especially great in applied research since decisions affecting individuals and public policies may be based on publications. Journals may therefore wish to know if an investigation has been started, and may wish to alert readers before an investigation (and appeal process) has concluded (e.g. by an expression of concern).

However, in many jurisdictions, research misconduct investigations and disciplinary hearings are considered confidential and institutions/employers may therefore feel unable to share details with journals. This approach may prevent journals from fulfilling their responsibilities to their readers, for example by publishing an expression of concern.

Various solutions to this problem were discussed at the CLUE meeting. One suggestion was for journals to require authors to disclose any allegations or proceedings and thus waive the confidentiality accorded by law within their contract with the journal. Another suggestion was that researchers’ employment contracts should specify that, in cases of suspected or proven misconduct, harm to research participants, or other circumstances affecting the validity of a research report, the employees’ usual right to confidentiality in disciplinary proceedings would be waived to allow the institution to communicate relevant details to the journal and other parties. The CLUE meeting participants recognised that such solutions might be hard for journals to enforce, or require changes in employment legislation, and therefore put them forward for discussion rather than as recommendations.

### Issue: Institutions sometimes do not share findings of misconduct investigations with journals that have published affected research and journals may be reluctant to publish informative retraction notices

Journals have a duty to avoid misleading their readers and therefore sometimes need to correct or retract published work that is incorrect or unreliable. Since problems can arise either inadvertently, from honest error, or from deliberate misconduct, retraction guidelines [14] recommend that the reason for a retraction should be clearly stated in the retraction notice including details of the affected findings and the type of problem detected.

This is important to ensure that honest researchers are not discouraged from alerting journals to problems with their work because of fears that a retraction will damage their career or be taken to imply that misconduct has occurred (when, in fact, such honesty and care for the research record should be praised [15]). Journals that have published affected work therefore need to receive details of misconduct investigations including clear information about all of the published articles (and submitted manuscripts) that are affected.

Being able to quote or cite an official report from an institution should facilitate the publication of clear and informative retractions (or corrections) since it reduces the journal’s risk of litigation. If a journal reports that University X has investigated the case and determined that a researcher has fabricated data this is a statement of fact and therefore unlikely to expose the journal to claims that it has published defamatory material.

Although, after misconduct has been found, institutions often require researchers to contact journals in which their work was published, we encourage institutions also to contact the journals directly. This direct communication between institution and journal allows relevant information to be shared and avoids situations in which researchers fail to contact affected journals, refuse to accept an investigation’s findings, or give a misleading account of the investigation to the journal. If an author tells a journal that the investigation was unfair or its finding was incorrect, this places the journal in a difficult position, but this problem may be avoided if the journal is allowed to see the full report of the investigation and can therefore verify whether it was properly conducted. We also recommend that institutions should be transparent about their processes for handling suspected misconduct or, at least be prepared to share information about such processes with journals, if requested.

#### Recommendations

Institutions should notify journals directly and release relevant sections of reports of misconduct investigations to all journals that have published research that was the subject of the investigation. Names may be redacted to ensure privacy.

Institutions should allow journals to quote from misconduct investigation reports or cite them in retraction statements and related publications (e.g. explanatory editorials or commentaries).

### Issue: Journals and institutions may be contacted by whistleblowers who conceal their identity, use pseudonyms or request anonymity

Institutions should have policies about whistleblower protection and about the handling of cases from anonymous whistleblowers. Such allegations should be considered on their merits rather than being dismissed automatically. Therefore, an individual’s refusal to reveal their name, use of a pseudonym, or request to remain anonymous, should not prevent either a journal or an institution from taking allegations seriously. However, both journals and institutions need reassurance that an allegation is well-founded and is not simply a personal vendetta and therefore they may request further details or information from the correspondent and, if this is not forthcoming, it is reasonable for journals not to raise the concern with the university or for an institution to decide not to proceed with an inquiry or full investigation. However, this is a matter of judgement for both journals and universities, so we recommend a flexible approach, depending on the seriousness of the alleged problem or behaviour and the plausibility of the evidence provided. Journals should not feel compelled to respond to vexatious complaints and editors may seek legal intervention for persistent or threatening behaviour.

#### Recommendation

Anonymous or pseudonymous allegations made to journals or institutions should be judged on their merit and not dismissed automatically.

### Issue: Journals and institutions may be asked about publications relating to research that took place many years ago

While investigation of historical research may pose more challenges than inquiries into more recent work, concerns should not be dismissed solely on the grounds that the research was done a long time ago. If plausible evidence of serious problems is raised, it should, ideally, be examined regardless of when the problems occurred. However, contacting authors and accessing original data may be increasingly problematic the more time has elapsed since the research was performed. It is therefore reasonable for journals and institutions to prioritise the investigation of recent over historical work.

Institutions should take responsibility for research performed under their auspices regardless of whether the researcher still works at that institution. Even if a researcher has moved to another institution, or has retired, the appropriate investigation should take place. This is another reason why institutions should have mechanisms for retaining data for at least 10 years and, ideally, permanently.

Investigations into the work of researchers who have died, are chronically incapacitated or have left research altogether, is especially difficult, however, institutions should make their best efforts to establish whether work is reliable, so that journals can determine whether readers should be alerted to concerns. Although probably a rare occurrence, this is another situation in which public data posting or effective retention of data by institutions would be beneficial and in which assessing the reliability of findings and reports needs to be separated from determining whether misconduct was committed.

### Issue: Concerns may be raised about research that involved several institutions

When research involves several institutions, there is usually one institution that takes a primary or coordinating role in relation to the funding. This primary institution should be the initial point of contact and take the lead in responding to concerns about the reliability of the research. Ideally, research agreements should specify this and also set out responsibilities for data deposition and retention [16].

The International Committee of Medical Journal Editors (ICMJE) states that authors should be accountable for answering questions about research and identifying which author was responsible for each aspect if questions arise [17]. We suggest extending this guidance so that authors are also expected to identify where each component of a project was done, and therefore which institution should be responsible for investigating any concerns about it.

### Issue: If a journal rejects an article about which either reviewers or editors have raised concerns about reliability, authors may simply submit it to another journal, perhaps after concealing problems more effectively

The COPE Code of Conduct notes that “Editors should not simply reject papers that raise concerns about possible misconduct. They are ethically obliged to pursue alleged cases.” [18] In other words, journals should seek explanations from authors even if they do not intend to accept their publication and should contact institutions, if required, regardless of publication status.

All research institutions should establish, promote, and incentivise a culture that encourages integrity of research and publications. This may involve rewarding mentorship and providing training on research integrity, peer review, and publication ethics. Such a commitment to integrity should also involve internal quality checks, but in many cases of research misconduct it is apparent that senior authors have not reviewed the data or thoroughly checked the validity and accuracy of the findings or the manuscript.

One suggestion made at the meeting was for each institution to maintain a repository of submitted manuscripts. Researchers affiliated to an institution would be expected to send a copy of all submissions to this repository. These would not be made public but the database could be used to check the history of a publication and document any changes made by authors (e.g. when submitting to a different journal after a rejection). Such a database of submitted manuscripts would be useful for institutional investigations and would permit assessment of all of a researcher’s work. To be workable this process would need to be straightforward and not excessively burdensome on researchers.

### Issue: Journals sometimes fail to respond to requests for correction or retraction from institutions or authors

Communication with a journal should normally be addressed to the editor, but if the editor does not respond, the publisher should be contacted. If a journal is owned by an academic society, the leaders of that society may also be used as a point of contact, or to raise concerns about the behaviour of the editor.

### Issue: Who should investigate if a peer reviewer is suspected of acting inappropriately?

Universities should recognize peer review as a legitimate part of research and academic activity and should encourage accountable and responsible behaviour from their researchers when they act as reviewers or editors [19]. However, even when peer review is viewed as part of general academic duties, the reviewer’s institution may not be equipped to investigate suspicions of reviewer misconduct since most of the relevant information will be held by the journal. In such situations, the journal may therefore have to initiate its own investigation, following the COPE flowchart about how to handle cases of suspected reviewer misconduct [11].

Evidence of serious misconduct by researchers acting as peer reviewers (e.g. stealing ideas or material from the articles they were invited to review) should be shared with their institution. Therefore journals should explain to reviewers that their identity might be disclosed to their institution in cases of suspected misconduct and that possible serious misconduct will be addressed by the institution.

### Issue: If a journal suspects that an author or peer reviewer has failed to disclose a relevant competing interest, should they refer this to the institution?

Readers, authors or reviewers sometimes suggest that relevant competing interests have not been disclosed during the review process or in a publication. If such allegations or concerns cannot be resolved (e.g. by publishing a correction if information has been omitted from a publication, or seeking additional peer review), the journal may consider contacting an institution. However, institutional responses vary. Some institutions maintain lists of researchers’ current interests and have policies about disclosure of competing interests. In such cases, it is appropriate for journals to raise concerns with the institution and to ask them for relevant information. However, not all institutions register such information, and, if they do not, they may be unable to respond to the journal’s enquiries. While failure to disclose a relevant interest is not always categorised as research misconduct, it is generally recognised to be poor practice and usually requires action by the journal (which will depend on the severity of the case).

#### Recommendation

Institutions and funders should be responsive to journal requests for information to ensure that peer reviewers’ and authors’ competing interests are properly disclosed.

## Recommendations on best practice

i. Institutions should have a research integrity officer (or office) and publish their contact details. National research integrity bodies (or other appropriate organizations, e.g. major funders) should keep a register of people responsible for research integrity at their country’s institutions, to enable journal editors (and others) to contact them. Where such lists are not available, journals should request corresponding authors to provide the name and contact details of their institution’s research integrity officer (or of an individual with responsibility for handling research integrity cases).
ii. Journals should develop criteria for determining whether, and what type of, information relating to the validity or reliability of research reports should be passed on to institutions. In addition to sharing any direct evidence of plagiarism, fabrication or falsification with institutions, journals should share reviewer or editor suspicions that work is “too good to be true” or of something being “not right”. Journals should not reveal the identity of peer reviewers or other people raising concerns (unless this is already published or the individuals have given permission for this disclosure). Anonymous or pseudonymous allegations to journals should be judged on their merit and not dismissed automatically.
iii. While journals should normally raise concerns with authors in the first instance, they should also have criteria to determine when the authors’ institution(s) should be contacted immediately without (or at the same time as) alerting the author(s).This would normally occur only in exceptional cases when journals have strong suspicions or clear evidence of substantive or significant falsification or fabrication of data.
iv. Research institutions and major funders should have systems to ensure that essential research data are retained for at least 10 years, and ideally permanently. Responsibility for data storage (e.g. for multicentre studies) should be defined in funding agreements.
v. Journals and publishers should retain peer review records for at least 10 years to enable the investigation of peer review manipulation or other inappropriate behaviour by authors or reviewers.
vi. Institutions should develop mechanisms for assessing the validity of research reports that are submitted to, or published by, academic journals; these processes should be independent from systems to determine whether misconduct has occurred.
vii. Institutions should publish their processes for conducting inquiries and investigating misconduct and should share information about such processes with journals, on request. Anonymous or pseudonymous allegations to institutions should be judged on their merit and not dismissed automatically.
viii. Institutions should notify journals directly and release relevant sections of reports of misconduct investigations to all journals that have published research that was the subject of the investigation. The report should clearly indicate which articles or manuscripts are affected. Names may be redacted to ensure privacy. Institutions should allow journals to quote from misconduct investigation reports or cite them in retraction statements and related publications (e.g. explanatory editorials or commentaries).
ix. Institutions and funders should respond to journal requests for information to ensure that peer reviewers’ and authors’ competing interests are properly disclosed.

## Proposals requiring further discussion

- Researcher employment contracts should indicate that the researcher’s name and relevant details of the affected research may be released to a journal or appropriate authority in cases of misconduct.
- Journals should require authors (as part of their publication contract) to disclose any allegations or proceedings relating to the submitted or published work.
- Institutions should maintain internal repositories of all submitted manuscripts so researchers’ work can be reviewed and changes to manuscripts identified, if needed.

